# CryoET structures of immature HIV Gag reveal a complete six-helix bundle and stabilizing small molecules distinct from IP6

**DOI:** 10.1101/2020.10.31.363382

**Authors:** Luiza Mendonça, Dapeng Sun, Jiying Ning, Jiwei Liu, Abhay Kotecha, Mateusz Olek, Thomas Frosio, Xiaofeng Fu, Benjamin A Himes, Alex B. Kleinpeter, Eric O. Freed, Jing Zhou, Christopher Aiken, Peijun Zhang

## Abstract

Gag is the major HIV-1 structural polyprotein precursor. The Gag SP1 domain with the last residues of CA have been hypothesized to form a six-helix bundle necessary for particle assembly, but this bundle has not been fully resolved. Here, we determined the structures of complete CA-SP1 six-helix bundle connecting to the NC domain, from both *in vitro* Gag assemblies and viral-like particles (VLPs) carrying a T8I mutation in SP1, to near-atomic resolutions using cryoET and subtomogram averaging. The structures revealed novel densities, however distinct from IP6, inside the six-helix bundle of Gag assemblies, stabilizing the immature lattice. Interestingly, the T8I mutation impaired proteolytic cleavage of Gag at both SP1 boundaries. Our findings signify the involvement of small molecules in immature Gag assembly and provide the structural basis for development of small molecule inhibitors that stabilize SP1 helix, thus interfere with PR-mediated virus maturation.

## Introduction

Gag is the structural precursor protein in HIV-1. It is a polyprotein synthesized from the unspliced viral mRNA, and is cleaved by the viral protease (PR) during the maturation stage of the replicative cycle into 6 proteins and peptides: matrix (MA), capsid (CA), spacer peptide 1 (SP1), nucleocapsid (NC), spacer peptide 2 (SP2) and p6^1^. Gag plays a major role during viral particle assembly, with each of its protein domains performing distinct functions. MA is responsible for membrane targeting. NC is responsible for RNA encapsidation. The CA and SP1 region multimerizes forming the immature viral lattice, comprised of interconnected hexamers. The Gag hexamer is, therefore, the assembly unit of the immature lattice. Proteolytic cleavage of Gag results in the transition of the immature Gag lattice to a mature capsid formed by the CA protein^2–4^.

The SP1 segment of Gag is 14 amino acids long and is delimited by the very first (SP1|NC) and the very last (CA|SP1) cleavage sites processed during HIV-1 maturation. SP1 plays a major role in the higher-order multimerization of Gag during viral assembly and is essential for viral infectivity^5–7^. Molecular dynamics and circular dichroism studies indicated that at high local concentrations, the stretch of residues comprising the last 8 amino acids of CA and the SP1 form a six-helix bundle^5,8^. It has been hypothesized that the amphipathic nature of this region would lead to association of six CA-SP1 molecules, thus burying the hydrophobic residues in the internal face of the bundle, thereby stabilizing the Gag hexamer^8^. An NMR study of this region indicated that it adopts an alpha-helical conformation in 30% TFE^9^, and at high protein concentrations in aqueous solutions^8^. Magic Angle Spinning (MAS) NMR of CA-SP1 tubular assemblies demonstrated that the SP1 peptide is highly dynamic^10^. The structures of immature CA lattice and the first 8 residues of SP1 were determined previously by cryoEM and X-ray crystallography^11,12^. Both structures show a six-helix bundle formed by a stretch of 16 residues, 8 from the CA C-terminal domain (CA_CTD_) and 8 from SP1. However, the rest of SP1 and its connection to NC could not be resolved in these structures.

Other factors are considered to be important for immature lattice formation and stability, in particular the small molecule inositol hexakisphosphate (IP6)^13–16^. The CA-SP1 six-helix bundle contains two rings of positive charge formed by 12 lysines, K158 and K227, in the CA_CTD_. The cryoEM and x-ray crystal structures indicate that all the lysine side chains point toward the center of the Gag hexamer, creating a strong positive potential^11,12^. IP6 was found to neutralize the positive charges at the center of the Gag hexamer, thus acting as a co-factor stabilizing the Gag hexamer. Other polyanionic molecules may also fulfil this role^17^, but they remain to be identified and characterized.

The SP1 region is thought to be the target of multiple maturation inhibitors (MIs), such as Bevirimat (BVM) and PF-46396 (PF-96)^18–23^. MIs block CA|SP1 cleavage, presumably by stabilizing the CA-SP1 region, and/or obstructing protease binding, thus impairing viral infectivity^11,12,24^. Natural polymorphisms as well as escape mutations in the CA_CTD_ and SP1 can render MIs ineffective^25,26^. Compensatory mutations selected during the passage of MI-dependent viral mutants were shown to phenocopy the effect of MIs. One such compensatory mutation is the single amino acid change, SP1 T8I, which is particularly interesting. It was selected independently during the propagation of PF-96 and BVM-dependent viruses in the absence of these MIs^22,25^. The T8I mutation was found to promote the assembly of the Gag lattice and stabilize the alpha-helical conformation of CA-SP1^19,24^.

Here we report the structures of two types of immature Gag assemblies bearing the T8I mutation: one from purified Gag assemblies made in bacterial cells where the Gag polyprotein was expressed; the other from purified VLPs produced in mammalian cells. We solved the structures of full-length CA-SP1 from both assemblies at 4.2 Å and 4.5 Å resolutions using cryo-electron tomography (cryoET) and subtomogram averaging (STA)^27^. Our maps show a fully extended six-helix bundle that connects the CA_CTD_ to the NC-RNA density layers and a possible organization of the NC-RNA. Intriguingly, novel densities, other than IP6, were identified to coordinate lysine residues in a manner similar to IP6, stabilizing CA-SP1 six-helix bundle. We further investigated the effect of T8I mutation on Gag processing, demonstrating that PR-mediated cleavage at both SP1 boundaries was impaired.

## Results

### CryoET and subtomogram averaging of GagT8I assemblies and VLPs

To resolve the structure of the full CA-SP1 six helix bundle, we utilized HIV-1 Gag proteins containing the T8I mutation at SP1 for stabilizing bundle assembly. SP1 T8I is a compensatory mutation that emerged in MI-dependent viruses subject to the selective pressure of replicating under sub-optimal concentrations of MIs. This mutation was shown to stabilize immature assembly^22,24,25^. We used two different Gag assemblies for structure analysis by cryoET: Gag_ΔMA_T8I protein assemblies expressed in *E. coli* (Fig. 1a and b) and GagT8I VLPs produced in HEK293T cells (Fig. 1a and d). The Gag_ΔMA_T8I construct lacks most of the globular domain of MA (residues 15 to 100) and p6 (Fig. 1a). Wild-type (WT) Gag_ΔMA_ is a widely used construct for HIV-1 Gag lattice investigation^11,13,28–30^. Upon induction in *E. coli*, the WT Gag_ΔMA_ protein was expressed, purified from bacterial lysates, and assembled *in vitro* into spherical particles bearing Gag immature lattices in the presence of oligonucleotides. The Gag_ΔMA_T8I protein, however, was not soluble when released from bacteria owing to its self-assembly into spherical particles inside the bacterial cells (Supplementary Fig. 1). We therefore purified Gag_ΔMA_T8I assemblies directly from lysed bacterial cells. The GagT8I VLPs were produced by transfection of a codon-optimized Gag expression vector into HEK293T cells and purified from the cell supernatant, similar to the WT Gag VLPs^31^.

**Figure 1.**
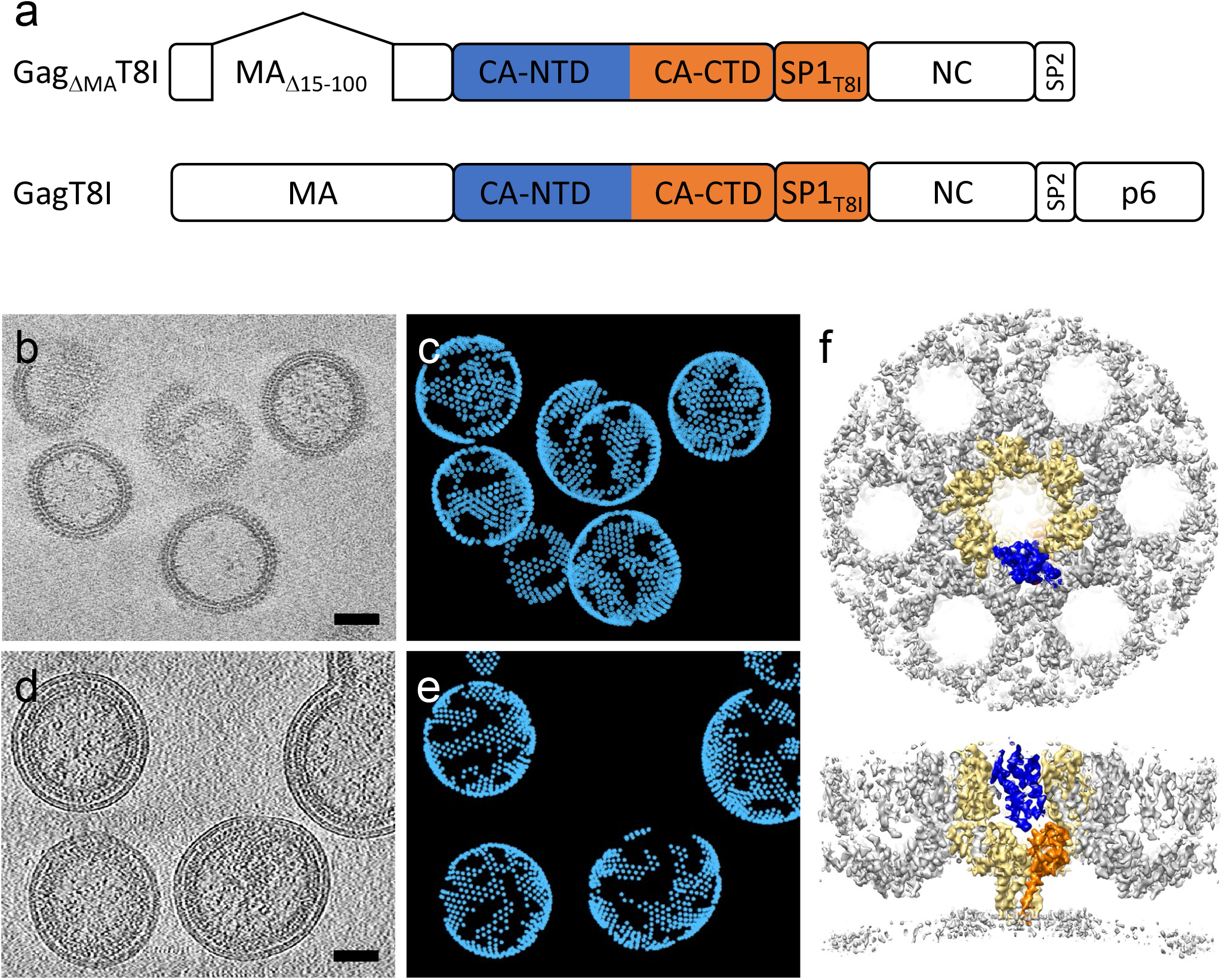
CryoET and subtomogram averaging of Gag T8I assemblies and VLPs. (a) Organization of the two constructs used for sample production: Gag_ΔMA_T8I, used for Gag assemblies made in *E. coli* cells; and GagT8I, used for VLP production in HEK293T cells. Both constructs bear the T8I mutation in the SP1 region (T371I in Gag sequence). (b-c) Purified Gag_ΔMA_T8I assemblies from *E. coli* cells are shown in a representative tomographic slice (b) and emClarity template matching result (c). (d-e) Purified GagT8I VLPs produced in HEK293T cells shown in a representative tomographic slice (d) and emClarity template matching result (e). (f) Subtomogram average of Gag_ΔMA_T8I assemblies in an extended lattice, shown in top and side views. The central Gag_ΔMA_T8I hexamer is highlighted in yellow and within which a monomer is highlighted in blue (CA_NTD_) and orange (CA_CTD_-SP1). Scale bars are 50 nm. Each point in (c) and (e) represents the position of a Gag hexamer.

A total of 90, 13 and 46 tilt series were collected for Gag_ΔMA_T8I assemblies in the absence and presence of IP6 and GagT8I VLPs, respectively. Details of data acquisition and image processing are summarized in Table 1, and data processing workflow is illustrated in Supplementary Figure 2. We created a cryoET Toolbox for the on-the-fly processing of cryoET data as they were acquired (Supplementary Fig. 2). emClarity, a software package developed in our group^32^, was used for subtomogram alignment and averaging. A low-pass filtered (25 Å) Gag hexamer density map (EMD-8403)^33^ was used for template-matching in emClarity, to automatically extract the position and initial orientation of the Gag hexamer in tomograms, as shown in Figure 1c and e. Upon iterative refinement (Supplementary Fig. 2), the final density maps of the Gag hexamer from Gag_ΔMA_T8I assemblies in the absence and presence of IP6 and GagT8I VLPs were obtained at 4.2 Å, 4.2 Å and 4.5 Å resolution, respectively (Supplementary Fig. 3c). The resulting maps confirm an immature Gag lattice from Gag_ΔMA_T8I assemblies and GagT8I VLPs, which is composed of interconnected hexamers formed by the CA N-terminal domain (CA_NTD_), CA_CTD_ and an extended six-helix bundle formed by CA_CTD_ and SP1 peptide (Fig. 1f).

**Table 1.**
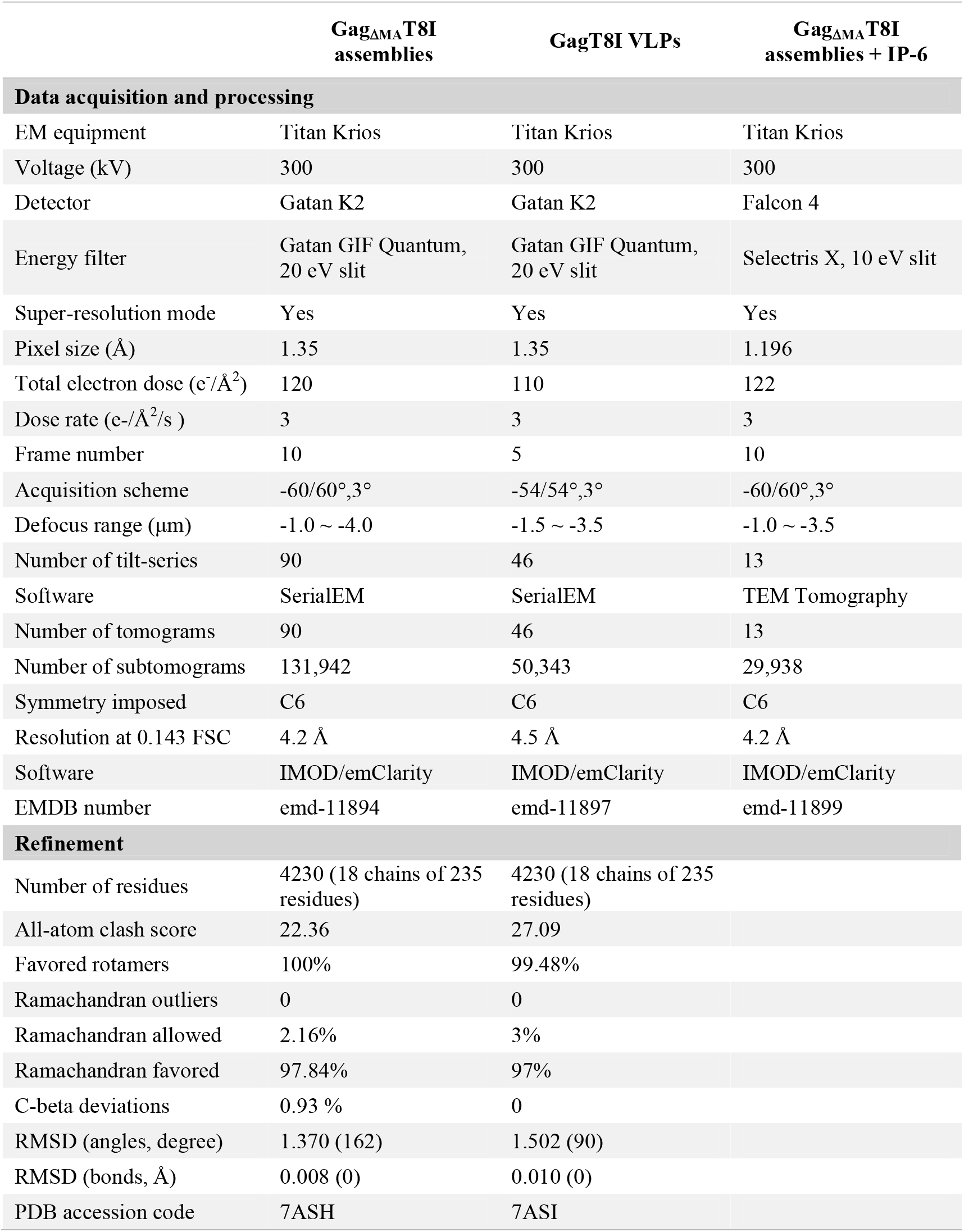
Cryo-EM data collection, refinement and validation statistics.

### GagT8I assembles into a full six-helix bundle further stabilized by small molecules

The subtomogram-averaged maps at 4.2 Å and 4.5 Å resolutions enabled refinement of atomic models of CA and SP1 (Fig. 2a-b). The refinement statistics are summarized in Table 1. Overall, the structures of Gag_ΔMA_T8I and GagT8I VLPs are very similar, with an RMSD of 0.577 Å (Fig. 2a-b). The complete CA-SP1 helix and the NC-RNA density layer which SP1 connects are clearly resolved in both maps (Fig. 2a-b), which were incomplete or missing from the previous CA and SP1 structures^11,12^. Nonetheless, interesting differences could be observed between the two. The GagT8I VLP subtomogram average shows a density that extends beyond the N-terminus of CA, where 5 residues in the junction region between MA and CA are resolved and modelled (Fig. 2c, bottom). This is because GagT8I VLPs have a complete MA domain that interacts with the lipid envelope; by contrast, in the Gag_ΔMA_T8I assemblies the globular domain of MA is missing and Gag is not constrained by a lipid envelope (Supplementary Fig. 4). The refined atomic model shows that the T8I mutation stabilizes the Gag hexamer by enhancing the hydrophobicity in one face of the amphipathic CA-SP1 helix (Supplementary Fig. 5). This confirms the hypothesis that the Gag hexamer is formed by the association and burial of the hydrophobic CA-SP1 face in the internal channel of the six-helix bundle^7,8^.

**Figure 2.**
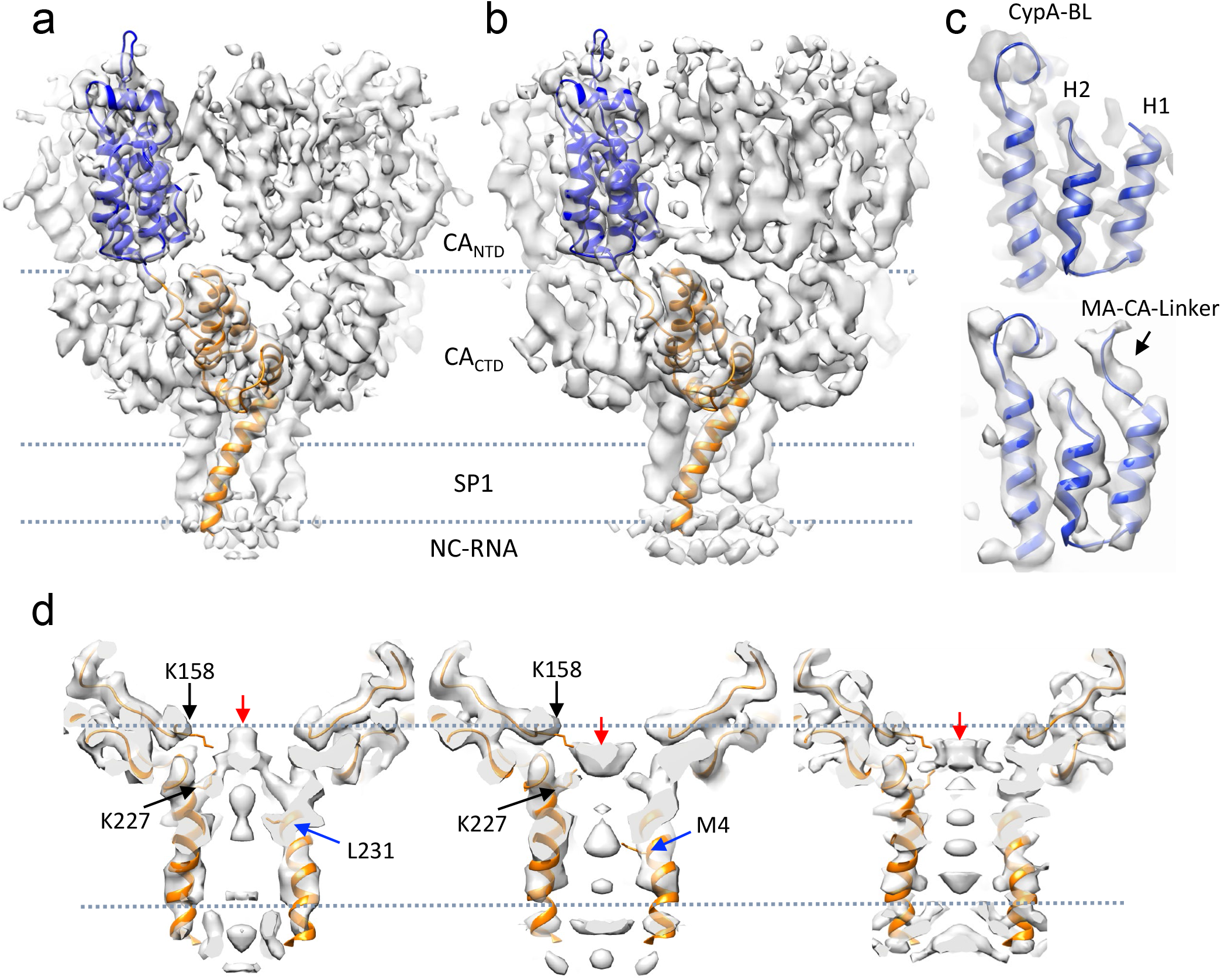
Structures of immature GagT8I CA-SP1 by subtomogram averaging. (a-b) The density maps of immature Gag CA-SP1 from Gag_ΔMA_T8I assemblies (a) and GagT8I VLPs (b), overlaid with the refined respective atomic models shown in blue (CA_NTD_) and orange (CA_CTD_-SP1). Domain regions are labeled. (c) Enlarged views of CypA-binding loop (CypA-BL) and N-terminal MA-CA linker in Gag_ΔMA_T8I assemblies (top) and GagT8I VLPs (bottom). (d) Detailed views of the internal densities within the six-helix bundle of Gag_ΔMA_T8I assemblies (left), GagT8I VLPs (middle) and Gag_ΔMA_T8I assemblies with 10μM IP6 (right). Black arrows point to the lysine residues from CA_CTD_, K158 and K227, forming two-rings to coordinate a central density (red arrows). Blue arrows point to residues CA L231 in Gag_ΔMA_T8I assemblies and SP1 M4 in GagT8I VLPs. Dashed lines mark the same height in both density maps.

Intriguingly, we observed a strong density inside the space formed by the six-helix bundle. This density is coordinated by two lysine rings formed by residues K158 and K227 near the bottom of the CA_CTD_ and above the bundle (Fig. 2d). A close inspection revealed that the profile and position of the internal density are different in the Gag_ΔMA_T8I and GagT8I VLPs maps (Fig. 2d, red arrows). For GagT8I VLPs produced in mammalian cells, this density is consistent with IP6, as suggested in previous studies^13,14^ (Fig. 2d, middle). IP6 is known to associate with the HIV-1 immature lattice and is present at cytoplasmic concentrations estimated at 20-50 nM in mammalian cells^34^. *E. coli* cells, however, lack the enzyme responsible for biosynthesis of the inositol ring and are thus thought to lack inositides^35^. Considering that Gag_ΔMA_T8I assemblies were produced in *E. coli* cells, it seems unlikely that the density in Gag_ΔMA_T8I map is IP6 (Fig. 2d, left). In fact, when the same Gag_ΔMA_T8I assembly is incubated with a buffer containing 10 μM IP6, this internal density reverts to a lower position closely resembling that seen in GagT8I VLPs (Fig. 2d. right). It is plausible that a small anionic molecule other than IP6 assists in the charge neutralization of the lysine rings and promoted assembly in *E. coli*-expressed Gag_ΔMA_T8I. Consistent with this, a similar density was also present at the same location in a previously published subtomogram average of bacterially expressed WT Gag_ΔMA_ assemblies (Fig. 3c, blue)^11^. Further studies are needed to identify such small molecules. Additional small molecules may also be involved in stabilization of the six-helix bundle and HIV-1 immature lattice, as another set of densities were observed inside the six-helix bundle below the lysine rings. These densities differ between the Gag_ΔMA_T8I and GagT8I VLP maps (Fig. 2d). In Gag_ΔMA_T8I, this density sits in the highly hydrophobic environment made by the L231 residue (L363 in Gag), whereas in GagT8I VLPs, this density is closer to the M4 residues (M367 in Gag). An exciting prospect from this study is that small molecules can be located and differentiated in subtomogram averages of *in situ* structures, though a better resolution would help in their identification.

**Figure 3.**
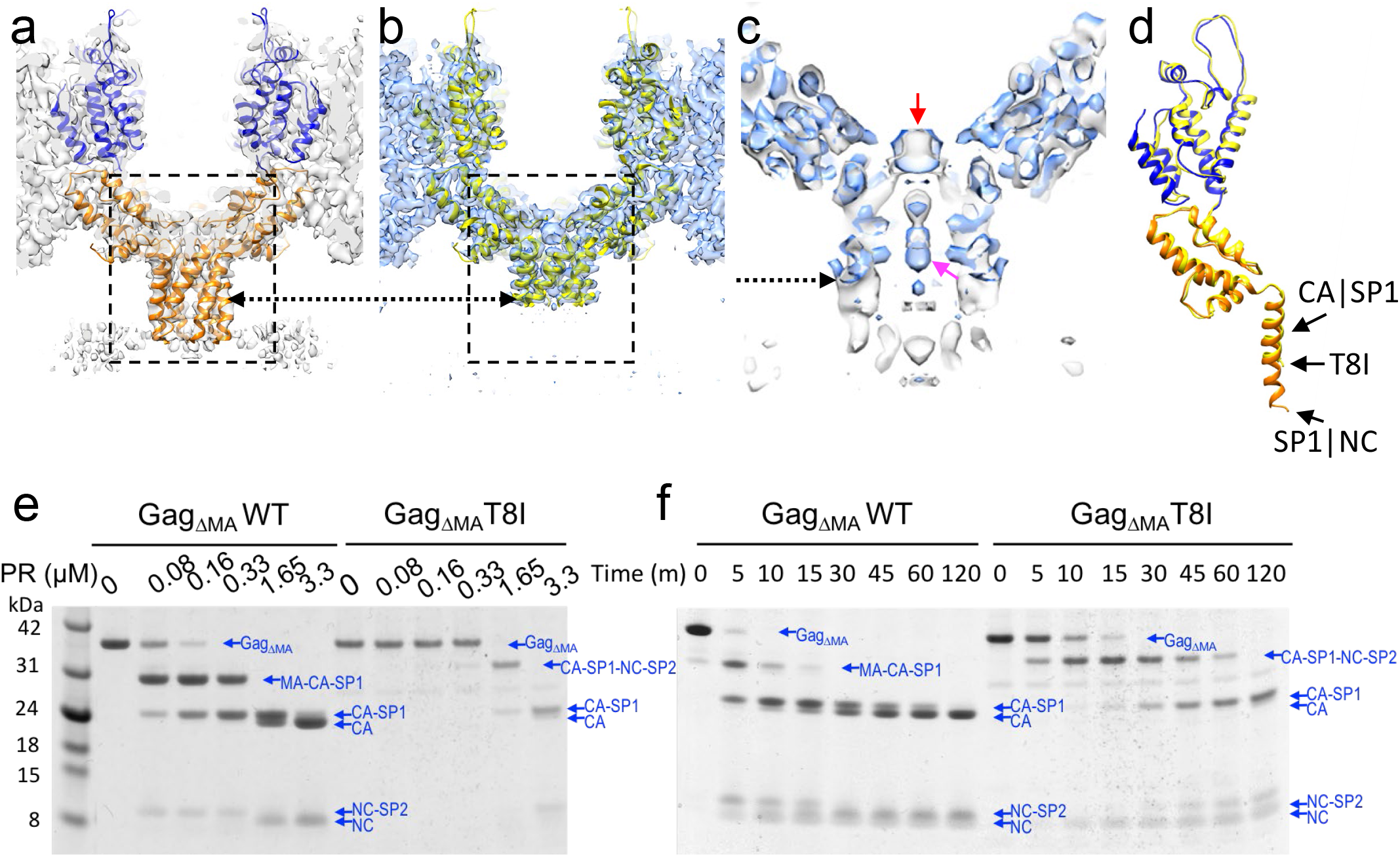
The T8I mutation stabilizes the six-helix bundle and impairs proteolytic processing of cleavage sites flanking SP1. (a-b) Comparison of the CA-SP1 hexamer in Gag_ΔMA_T8I (a) and Gag_ΔMA_WT (EMD-3782) (b), shown in a side-view. The density map is overlaid with the respective atomic model. The dashed arrows point to the position of T8I in both maps. (c) An enlarged view of density overlay from the dashed box region in a and b. The Gag_ΔMA_T8I density is in grey and the Gag_ΔMA_WT density is in blue. The red arrow points to the same central density shown in Figure 2d. The pink arrow points to BVM density in EMD-3782. (d) Overlay of the refined atomic models from Gag_ΔMA_T8I and Gag_ΔMA_WT (PDB 5L93). The arrow points to the location of the T8I mutation and CA|SP1 and SP1|NC cleavage sites. (e-f) *In-vitro* PR cleavage assays. Gag_ΔMA_ WT and Gag_ΔMA_T8I assemblies were incubated with recombinant HIV-1 PR at the indicated concentration (e) and for the indicated period of time (f), separated by SDS-PAGE and stained with Coomassie Blue. The protein bands are labelled.

### The T8I mutation reduces SP1 dynamics and impairs proteolytic cleavage

In the previous WT Gag_ΔMA_ map (EMD-3782) and its corresponding atomic model (PDB 5L93), the SP1 region after T8 was not resolved^11^. The SP1 helix in the Gag_ΔMA_T8I map is complete and extends to the SP1 and NC interface (Fig. 3a-c), therefore allowing model building of the C-terminal portion of SP1, from T8I to M14 (Fig. 3d). The T8I mutation was shown to quench the dynamics of SP1 by MAS NMR^24^, and potentially stabilize a continuous helical conformation. It is worth noting that the WT Gag_ΔMA_ map (EMD-3782) was determined in the presence of BVM. The density corresponding to BVM is visible in the WT Gag_ΔMA_ map (Fig. 3c, pink arrow), but not in Gag_ΔMA_ T8I map (Fig. 2d, left panel). Yet, the observed unknown densities coordinated by K158 and K227 overlapped well in both WT Gag_ΔMA_ and Gag_ΔMA_T8I maps (Fig. 3c, red arrow), even though the two proteins were expressed and assembled independently in different manners in the absence of IP6.

The SP1 peptide is flanked by the first (SP1|NC) and the last (CA|SP1) cleavage sites in the HIV-1 maturation cascade. If the T8I mutation stabilizes the extended six-helix bundle, this may affect PR processing of these cleavage sites. We thus tested the effect of the T8I mutation on PR-mediated cleavage of Gag assemblies using *in vitro* cleavage assays (Fig. 3e-f)^36^. We first analysed the concentration-dependent processing of WT Gag_ΔMA_ and Gag_ΔMA_T8I by PR. WT Gag_ΔMA_ processing showed the established ordering of cleavage as the PR concentration increased, with the first major cleavage products MA-CA-SP1 and NC-SP2 (cleaving at SP1|NC), followed by CA-SP1 (cleaving at MA|CA), and completed with CA and NC (Fig. 3e). In contrast, not only was a much higher PR concentration required to initiate processing of Gag_ΔMA_T8I, but also the order of cleavage was altered (Fig. 3e). The first cleavage occurred at MA|CA site, suggesting an impaired or delayed cleavage at the SP1|NC site. Furthermore, the CA|SP1 site was also protected from cleavage, such that very little CA was produced. The same trend was observed when we investigated the kinetics of PR processing *in vitro* (Fig. 3f). At the end point of PR processing (120 minutes), CA-SP1 was the major final product for Gag_ΔMA_T8I, instead of CA. Therefore, cleavage at both proteolytic sites flanking SP1 are impaired by the T8I mutation, and/or the small molecules, which supports the formation of a stable and extended SP1 helix.

### The NC-RNA layer is defined but variable in the SP1-NC domain organization

We next sought to investigate how the SP1 six-helix bundle connects to the NC-RNA region, which has been intractable previously. As shown in Fig. 4a, three layers of densities were observed in tomographic slices of the Gag_ΔMA_T8I particles. Radial density profiles from three different particles of similar size displayed three distinct peaks corresponding to the CA_NTD_, CA_CTD_ and NC-RNA density layers (Fig. 4a)^37^. While CA_NTD_, CA_CTD_ and SP1 are rigidly connected as seen in the Gag_ΔMA_T8I map, NC-RNA is flexibly linked to SP1, resulting in a smeared density in Gag_ΔMA_T8I subtomogram averages (Fig. 3a).

**Figure 4.**
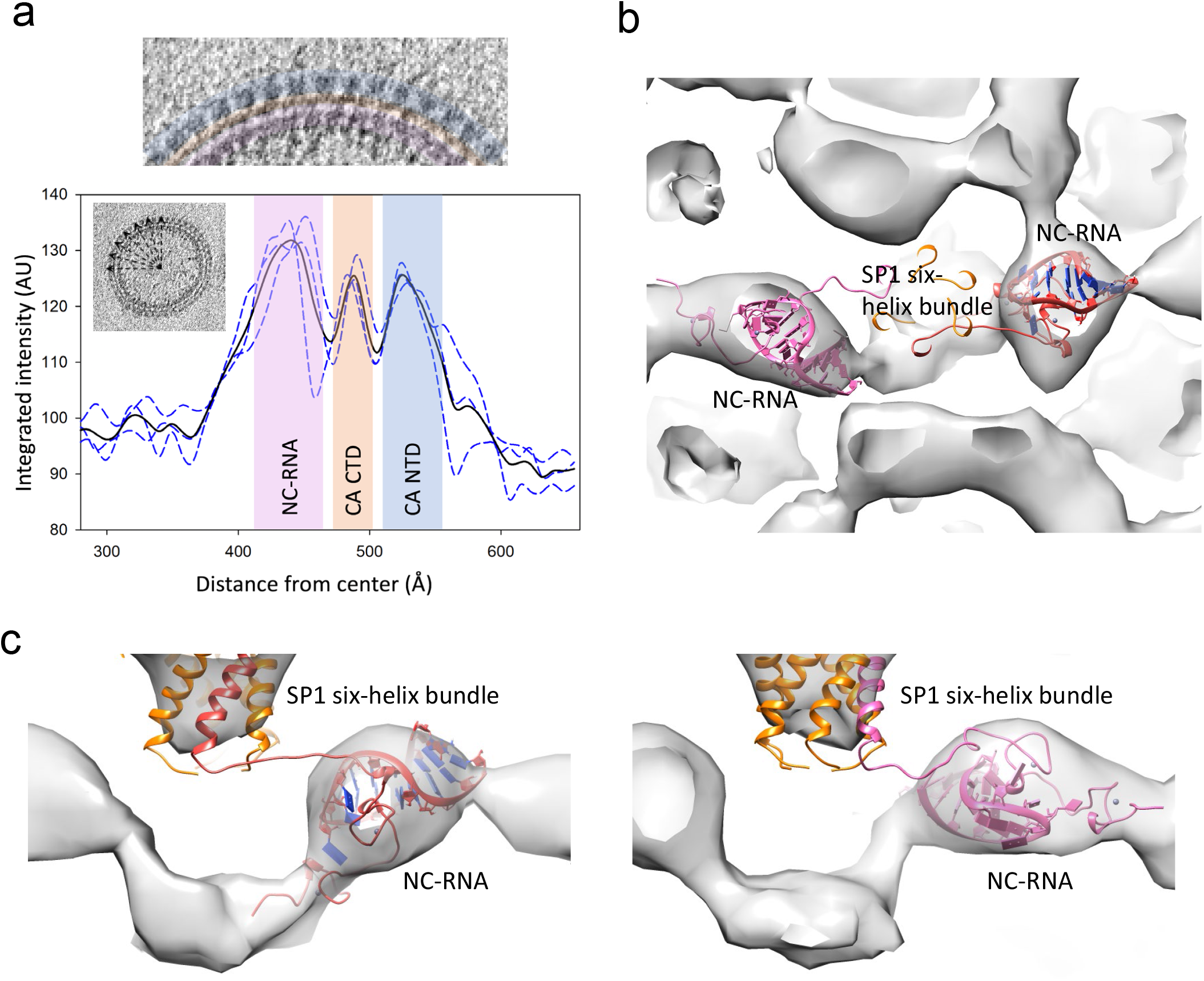
Characterization of the NC-RNA density layer. (a) Radial density profile of the Gag lattice. Top panel shows a tomographic slice of Gag_ΔMA_T8I lattice domain organization where CA_NTD_, CA_CTD_ and NC-RNA density layers are coloured in blue, orange and pink, respectively. Graph shows the radial density profiles of Gag_ΔMA_T8I particles of similar size (n=3). Black line indicates the average intensity while blue lines depict the individual particle profiles. Density profile peaks corresponding to CA_NTD_, CA_CTD_ and NC-RNA are shaded with the same colour scheme as in the top panel. (b) Top view from density map of six-helix bundle and the NC-RNA layer, overlaid with two plausible NC-RNA (ѱ SL2 loop) models, red and pink, fitted into the density, leaving the remaining NC-RNA density unmodeled. (c-d) Enlarged side view from density map in (b).

To gain structural information on the SP1-NC-RNA region, we carried out localized 3D alignment and classification, focusing on the SP1-NC-RNA region, using PCA and k-means subtomogram classification tools implemented in emClarity without imposing any symmetry (Supplementary Fig. 6a). Among 32 resulting 3D classes, we identified multiple classes that seemed to share common organization of the NC-RNA layer (Supplementary Fig. 6b, red boxes) and averaged 6747 particles from these classes to generate a density map for the SP1-NC-RNA region (Fig. 4b-c). A composite molecule of CA_CTD_-SP1-NC was created based on the refined atomic model CA-SP1 and the published NC-RNA model (PDB 1F6U)^38^, which was then used separately for rigid body fitting of the NC and the RNA domains in the density map. We performed this for each CA monomer in the model, resulting in two plausible models of SP1-NC-RNA organization (Fig. 4b-c). To bridge the space between the six-helix bundle density and the NC-RNA density layer, it was necessary to model the N-terminus of NC as a coil, up to residue Asn8. In both models, the NC-RNA complex is at the density layer below and surrounding the six-helix bundle (Fig. 4c). As HIV-1 NC is very flexible and the viral RNA can adopt many conformations, it is plausible that there exist multiple modes of SP1 to NC connection, including the two modes described here. Taken together, a stretched coil conformation at the SP1 and NC junction and/or dynamic unfolding of this region could facilitate PR processing. As such, the helix stabilization imparted by the T8I mutation may explain the observed delays in SP1|NC cleavage.

## Discussion

HIV-1 Gag is a multidomain and multifunctional protein responsible for membrane targeting, RNA encapsidation, particle assembly and budding. The CA and SP1 domains of Gag form the viral immature hexagonal lattice. Residues that make up the CA-SP1 helix are remarkably conserved and mutations in this region lead to loss of infectivity and/or assembly defects^8,19,39,40^.

The SP1-T8I mutation is a stabilizing compensatory mutation selected for during the passage of MI-dependent virus. Our refined atomic model shows that this mutation increases the hydrophobicity at the inner face of the six-helix bundle, thus stabilizing it (Supplementary Fig. 5). It has been shown that IP6 is also involved in Gag hexamer stabilization by neutralizing positive charges of lysine residues at the base of the CA_CTD_ ^13–16^. Here, we show that small molecules other than IP6 can neutralize charged lysine rings to stabilize the immature lattice, demonstrating the potential of subtomogram averaging to localize and differentiate small molecules in *in situ* structures, and possibility of developing small molecules as MIs..

HIV-1 maturation is a highly regulated process. The sequence of cleavage events is thought to be determined by the affinity constants and processing kinetics between the different cleavage sites in Gag and HIV-1 PR^41–43^. Events that alter the kinetics or sequence of maturation invariably lead to loss of infectivity^19,24,44,45^. T8I mutation was acquired as a second-site mutation that restores assembly and maturation to the BVM- and PF-96-dependent mutants, but the T8I single mutant exhibited a maturation defect with inefficient cleavage at CA-SP1. The inference is that the original resistant mutants exhibited unstable Gag lattices in the absence of the compound, which stabilizes the lattice, and T8I rescues the dependence on the compound by stabilizing the lattice, which becomes hyper-stable in the absence of a destabilizing mutation. We show that the T8I mutation alters the cleavage sequence in HIV-1 maturation and impairs processing of both SP1 boundaries, suggesting that a finely tuned SP1 stability is required to balance between lattice assembly, proper maturation and infectivity.

SP1 is critically important as one of the major targets for therapeutic drugs against HIV-1 and AIDS. The structure of full-length CA-SP1 in immature Gag assemblies derived from this study provides a blueprint for future development of small molecule inhibitors that can lock SP1 in a stable helical conformation and thus interfere with PR-mediated processing and virus maturation, and block HIV-1 infection.

## Materials and Methods

### Particle production and purification

The Gag_ΔMA_T8I mutant was constructed from pET2 PRR400 by site-direct mutagenesis. Protein was expressed in *E. coli*, Rosetta 2 (DE3), cultured in Luria-Bertani medium, and induced with 0.5 mM isopropylthiogalactoside at 23 °C for 16 hrs. Gag_ΔMA_T8I assemblies were formed inside of bacterial cells and purified directly from cell lysate. The cell pellets were collected and resuspended in lysis buffer (25 mM Tris, pH 7.5, 0.5 M NaCl) and broken with a microfluidizer. Subsequently, the lysate was centrifuged at 5000g for 10min to remove cell debris. The supernatant was collected and subjected to sucrose gradient centrifugation (30-70%) with a 15% sucrose cushion on the top. Gradient was spun at 210,000g for 18 hrs. Particles were collected from the pellet and resuspended in lysis buffer with and without 10 μM IP6 and used for CryoEM.

The GagT8I mutant VLPs were produced from HEK293T cells transfected with a codon-optimized full-length Gag expression plasmid (pCMV-Gag-opt)^46^. The culture supernatants were harvested approximately 40 hours after transfection, filtered through a 0.45-micron pore-sized filter, and pelleted through a cushion of 20% sucrose (wt/vol) in STE buffer (20 mM Tris-HCl pH 7.4, 100 mM NaCl, 1 mM EDTA). The VLPs were gently resuspended in STE buffer and frozen at −80°C.

### CryoEM sample preparation

A homemade manual plunger was used for cryoEM grid preparation. Quantifoil R2/2 copper grids were freshly glow discharged at 30 mA for 30 seconds. For Gag_ΔMA_T8I assemblies, 2 μl sample was applied to the carbon-side of the grid, followed by 1 μl 10 nm fiducial gold bead solution, and blotted from the backside since samples are very fragile and sensitive to blotting paper. 1 μl sample buffer was then added on the grid backside to facilitate the filter paper blotting from the backside. For GagT8I VLPs, 1 μl 10 nm fiducial gold beads were applied to the grid first, followed by 1~2 μl sample, and blotted from the frontside. The grid after blotting was then quickly immersed into liquid ethane cooled by liquid nitrogen. Frozen grids were stored in liquid nitrogen until data collection.

### CryoET data acquisition

Data for Gag_ΔMA_T8I assemblies and GagT8I VLPs were collected at the Electron Bio-Imaging Centre at Diamond Light Source (eBIC-DLS) in the United Kingdom. Tilt-series were acquired on a Gatan K2-Summit detector in super-resolution mode using a Thermo Fisher Scientific KriosG3i microscope operating at 300 kV equipped with a Gatan energy filter (slit width 20 eV; GIF Quantum LS, Gatan). Tilt-series were recorded at a nominal magnification of 105,000×, corresponding to a calibrated pixel size of 1.35 Å. A dose-symmetric scheme^47^ was used to collect tilt-series from −60° to 60° or −54° to 54° at a step size of 3° using SerialEM software^48^. At each tilt, a movie stack consisting of 5-10 frames was recorded with a set dose rate of 3-6 e^−^/Å^2^/sec. Tilt-series were collected at a range of nominal defocus between −1.0 and −4.0 μm and a target total dose of 110-120 e^−^/Å^2^ was applied over the entire series. For Gag_ΔMA_T8I with 10 uM IP6 sample, data were collected at the Thermo Fisher Scientific RnD division in Eindhoven, The Netherlands. Tilt-series were acquired on a Falcon4 detector in EER (electron event representation)^49,50^ format using a Thermo Fisher Scientific KriosG4 operating at 300 kV equipped with a CFEG and SelectrisX energy filter (slit width 10 eV; Thermo Fisher Scientific). Tilt-series were recorded at a nominal magnification of 105,000×, corresponding to a calibrated pixel size of 1.196 Å. A dose-symmetric scheme was used to collect tilt-series from −60° to 60° at a step size of 3° using TEM Tomography software. The CFEG was automatically flashed every 8hrs. At each tilt, a movie stack consisting of 217 EER frames was recorded with a dose rate of 4.6 e^−^/px/sec and a total dose of 2.98 e^−^/A^2^ per tilt. Tilt-series were collected at a nominal defocus range between −-1.0 and −4.0 μm and a target total dose of 122 e^−^/Å^2^ was applied over the entire tilt-series. Further details are given in Table 1.

### CryoET image processing

CryoET Toolbox (https://github.com/ffyr2w/cet_toolbox) was used for on-the-fly pre-processing of K2 datasets (freely available). In summary, movie frames were Fourier-cropped to a final pixel size of 1.35 Å and motion corrected by averaging frames for each tilt using program “alignframes” implemented in the IMOD package^51^ or MotionCor2^52^. For the Falcon4 data, 30 EER frames were grouped to create 7 dose fractions from 210 frames (last 7 frames were discarded) and motion corrected using Relion3.1^53^. Tilt series were aligned using the default parameters in IMOD version 4.10.22 with the eTomo interface^51^, using gold-fiducial markers. The alignment parameters including the projection transformations, local alignments, and fitted tilt angles were then passed to emClarity^32^ framework. Sub-tomogram alignment and averaging were carried out in emClarity following the published protocol^32^. Briefly, the workflow was as follows: Particles were picked from 6× binned non-CTF-corrected tomograms by emClarity template matching function using EMD-8403^33^ low-pass filtered to 25 Å as template. The template matching results were cleaned automatically on basis of geometrical restraints using ‘removeNeighbours’ function implemented in emClarity. Only particles that had at least 3 neighbours within 100 Å and oriented within 20° were retained. Subtomograms at the air-water interface were manually discarded using IMOD^51^. Following template matching, the data set was randomly split into two groups based on tomogram of origin, rather than randomly by sub-tomograms, which were processed independently for all subsequent steps. The initial positions and orientations of the first cycle averaging come from template matching results. C6 symmetry was applied throughout all sub-tomogram averaging procedures. The 3D alignment procedures were carried out gradually with binning of 5, 4, 3, 2 and 1. At each binning, duplicate particles were removed and the tilt-series geometry was refined using the positions of subtomograms as fiducial markers (TomoCPR). The Fourier Shell Correlation was calculated by the gold-standard method from even and odd data sets. Density maps were visualized in Chimera^54^ or PyMol (Schrödinger, Inc.). A diagram for the data processing is presented in Supplementary Fig. 2.

### Model building and refinement

An atomic model was built and refined against the Gag_ΔMA_T8I density map. The crystal structure (PDB 4XFX)^55^ and cryoEM structure (PDB 5L93)^11^ of HIV-1 CA were used as an initial reference. At 4.2 Å resolution, positive and bulky side chains are clearly visible enabling reasonable positioning of the residues in the atomic models. Rotamers were generally not refined, unless there was good evidence for a different rotamer in our density map. For the SP1 region, a composite model was made by joining the SP1 residues from PDB 1U57^9^ to the 5L93 model (lacking the last 6 SP1 residues) to serve as initial reference for the model refinement.

The model refinement procedure involved iterative manual refinement in Coot^56^ followed by 3 rounds of Phenix cryoEM real-space refinement^57^. After each round of real-space refinement, Ramachandran and rotamer outliers were manually refined in Coot, and another round of real-space refinement was performed until good model statistics were achieved. In order to ensure reasonable modelling in the SP1 region, the density map of Gag_ΔMA_T8I was sharpened with a B factor of 120 in combination with a 1% FDR confidence map^58^ and a locally sharpened B-factor map^59^ for the manual refinement in Coot.

### Subtomogram classification and modelling of the NC-RNA layer

For focused alignment and classification at the SP1-NC junction, a 49 × 49 × 47 Å mask was applied to the region that comprised the CA-CTD, the six-helix bundle and the NC-RNA layer. PCA and k-means classification implemented in emClarity were used to identify common organization features in NC-RNA layer in the subtomograms. No symmetry was imposed and tomograms were binned by a factor of 3 (equivalent pixel size 4.05 Å). Subtomograms were classified into 32 classes and among those 3 major classes were combined, comprising 6747 subvolumes.

A composite model was built in Coot^56^ using the refined Gag_ΔMA_T8I model and the published NC-ΨRNA_SL2_ model (PDB 1F6U)^38^. UCSF Chimera^54^ was used for rigid body fitting of the NC and RNA region of the composite model into the NC-RNA density map. This was repeated for each CA monomer. Bond lengths were regularized in Coot by using the Regularize Zone from residue M14 in SP1 to residue Q9 in NC.

### *In vitro* cleavage of Gag assemblies

PR digestion experiments with recombinant WT and T8I assemblies were performed by the addition of various concentrations of PR to the Gag WT assembly mixture and T8I cell lysate which was pre-treated and diluted to 2mg/ml, and incubated at 37°C for 2hrs. For kinetic analysis of Gag WT and T8I cleavage, 3.3μM of HIV-1 PR was added to the Gag assembly mixture and incubated at 37°C, at different time points, 4ul of the digestion reaction mixture was taken out and mixed with NuPAGE® LDS Sample Buffer (Invitrogen) to stop the reaction, and then subjected to NuPAGE Novex 4–12% Bis-Tris gel (Invitrogen) for cleavage products analysis and visualized by Coomassie blue staining.

## Data availability

The cryoEM density maps CA-SP1 hexamer from Gag_ΔMA_T8I assemblies in the absence and presence of IP6 and GagT8I VLPs were deposited in the EMDB under accession code EMD-11894, EMD-11899, and EMD-11897. The refined models were deposited in PDB under accession codes 7ASH and 7ASI, respectively.

## Acknowledgements

We thank Dr. Teresa Brosenitsch for critical reading of the manuscript. We are grateful to Drs Alistair Siebert, Kyle Dent, Andy Howe and Lingbo Yu for technical support. We thank Reint Boer Iwema for support with TEM Tomography software. We acknowledge Diamond for access and support of the CryoEM facilities at the UK national electron bio-imaging centre (eBIC, proposal NR18477, NR21005 and NT21004), funded by the Wellcome Trust, MRC and BBSRC. We also thank Thermo Fisher Scientific for microscope access. The computational aspects of this research were supported by the Wellcome Trust Core Award Grant Number 203141/Z/16/Z and the NIHR Oxford BRC. This work was supported by the National Institutes of Health P50 grant AI150481 (P.Z.), the UK Wellcome Trust Investigator Award 206422/Z/17/Z (P.Z.), and the UK Biotechnology and Biological Sciences Research Council grant BB/S003339/1 (P.Z.).

## Author contributions

P.Z. conceived the research and designed the experiments. J.N. purified Gag_ΔMA_T8I assemblies and performed *in vitro* protease cleavage experiments. J.Z. and C.A. purified GagT8I VLPs. X.F prepared cryoEM samples. J.L. and A.K. collected cryoET data. T.F. developed CryoET Toolbox used for pre-processing. L.M., D.S. and B.H. performed subtomogram averaging using emClarity. L.M., and M.O. refined the structural models. A.B.K. performed functional assays. L.M., E.O.F., D.S. and P.Z. analyzed data and wrote the paper with support from all the authors.

