## Supplemental figures for "CryoET structures of immature HIV Gag reveal a complete six-helix bundle and stabilizing small molecules distinct from IP6"

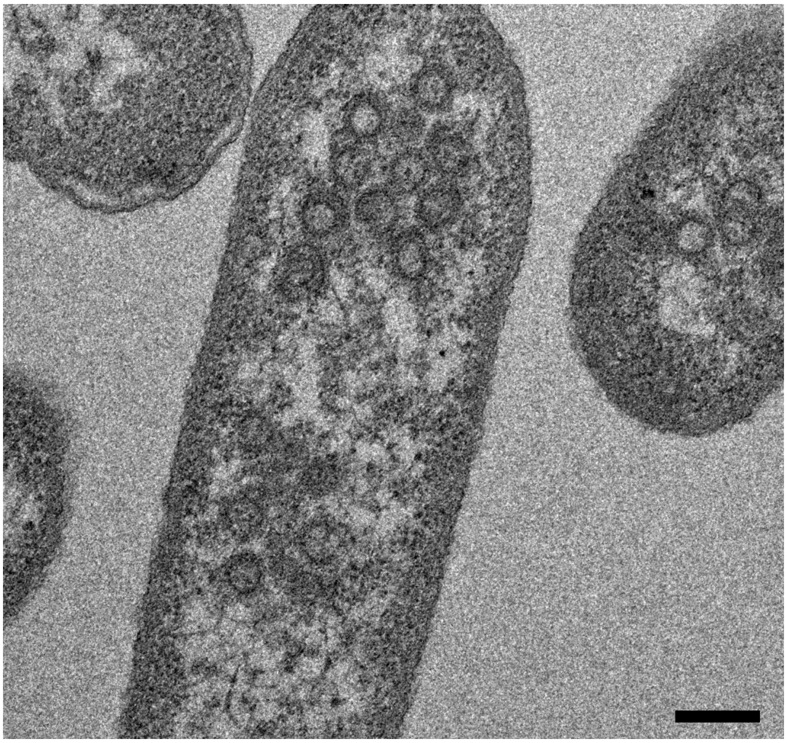

**Supplementary Figure 1 | Gag$_{\Delta MA}$T8I protein spontaneously assembles in *E. coli* when expressed.** Thin-section EM depicting spherical assemblies of Gag$_{\Delta MA}$T8I. Scale bar is 200 nm.

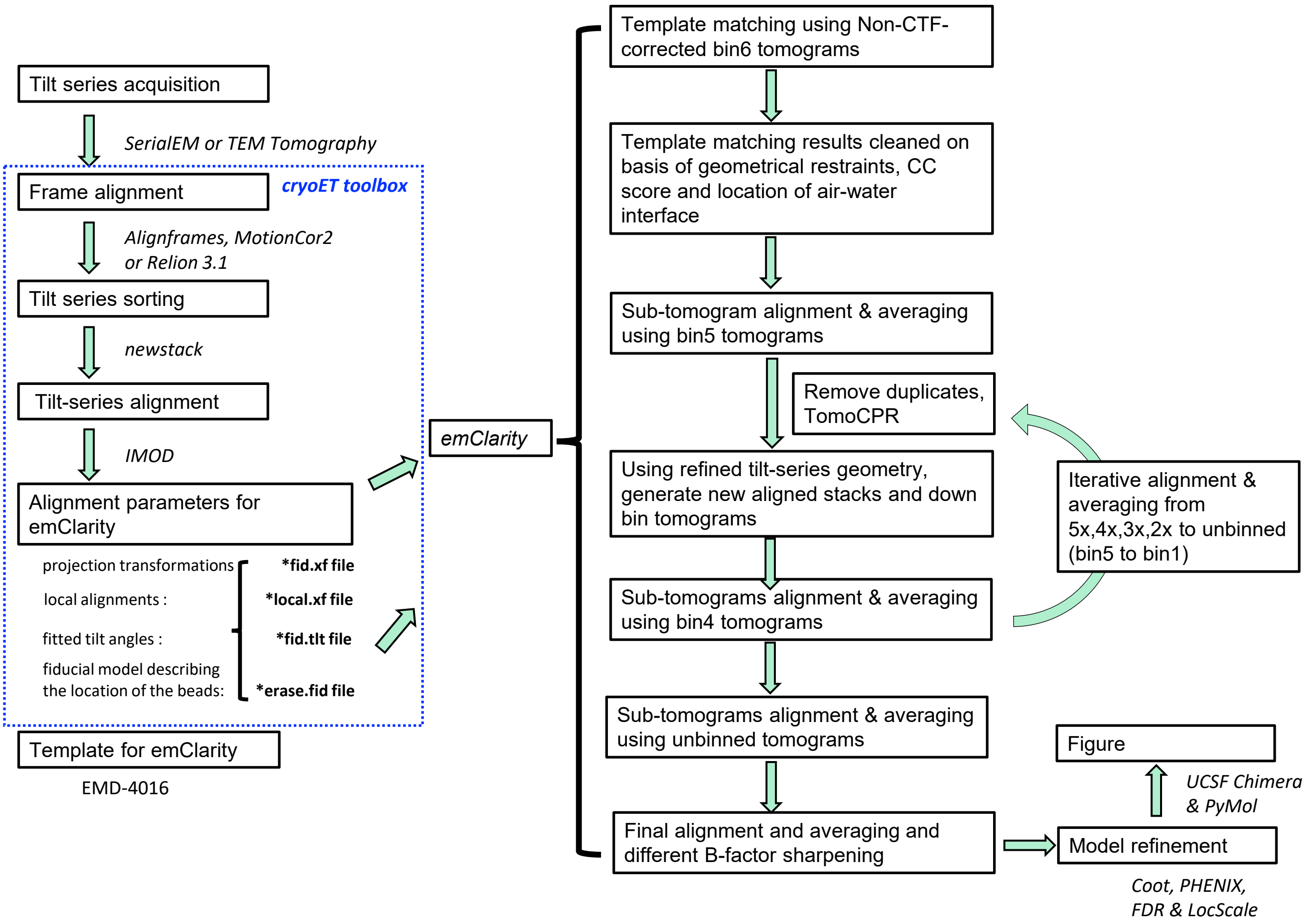


**Supplementary Figure 2 | Workflow data processing model refinement.** K2 Tilt-series were acquired using SerialEM [1]. Pre-processing was done on-the-fly using CryoET Toolbox (https://github.com/ffyr2w/cet_toolbox). In summary, movie frames were motion corrected and fourier-cropped by a factor of 2 using IMOD "alignframes" [2] or MotionCor2 [3]. Falcon 4 tilt-series were acquired with TEM Tomography and motion corrected using Relion 3.1 [4]. Tilt series were aligned using fiducial markers in eTomo[2]. The tilt-series alignment files were then passed to emClarity for sub-tomogram averaging [5]. Particles were picked from 6x binned non-CTF-corrected tomograms by emClarity template matching function using EMD-4016 [6] low-pass filtered to 25 Å as template. The template matching results were cleaned automatically on basis of geometrical restraints using 'removeNeighbours' in emClarity and subtomograms at the air-water interface were manually discarded using IMOD [2]. Following template matching, the data set was randomly split into two groups, which were processed independently for all subsequent steps. The 3D alignment was carried out with decreasing binning from 5 to 1. At each binning, duplicate particles were removed and the tilt-series geometry was refined using the positions of subtomograms as fiducial markers (TomoCPR). C6 symmetry was applied throughout the procedure. The Fourier Shell Correlation was calculated by the gold-standard method from even and odd data sets. Density maps were visualized in Chimera[7] or PyMol (Schrödinger, Inc.). Model refinement was carried out in COOT [8] and Phenix [9] aided by False-discovery Rate [10] and LocScale [11] maps.

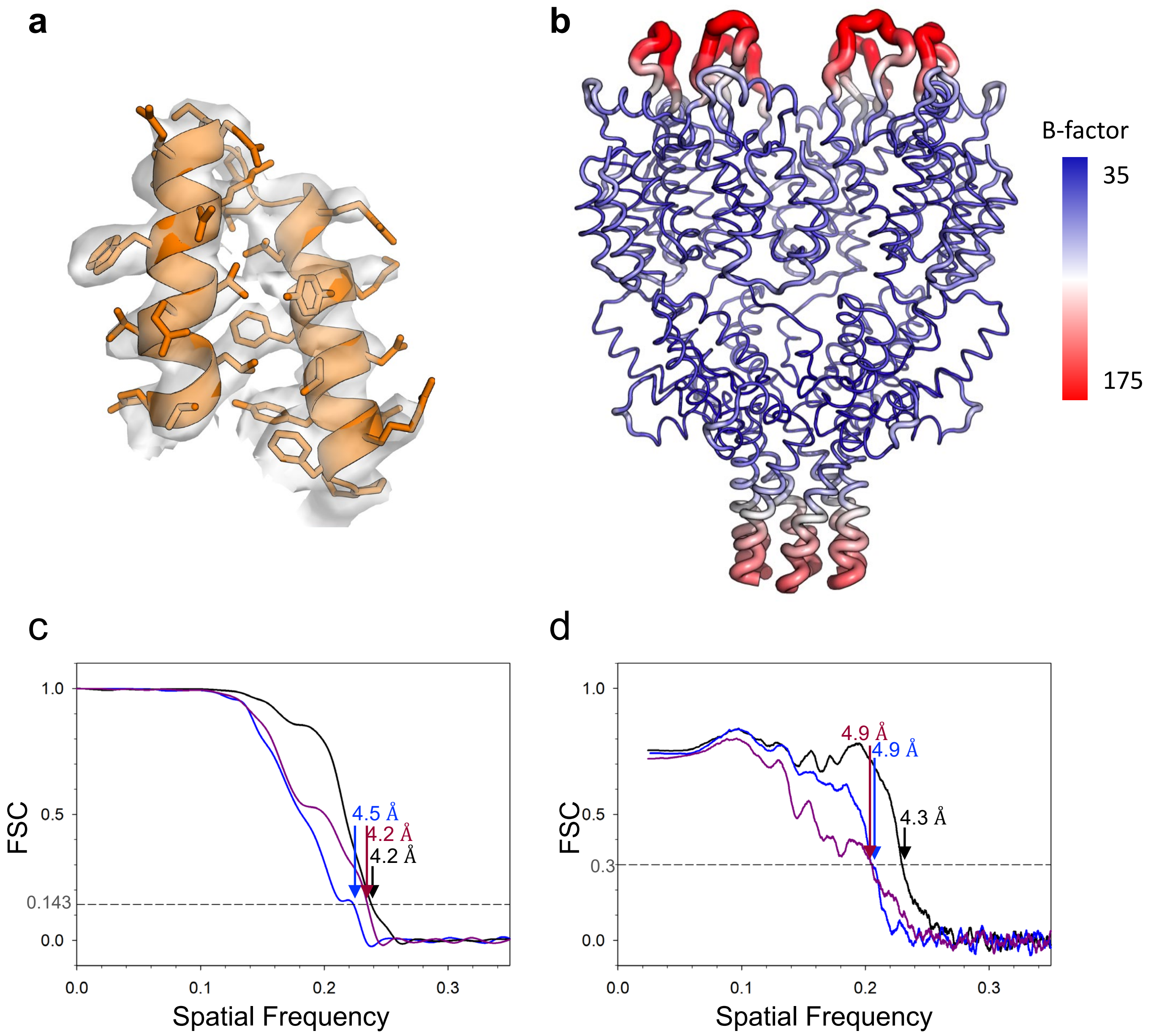


**Supplementary Figure 3 | The refinement of immature CA-SP1 models from GagT8I .** (a) Representative isosurface of $Gag_{\Delta MA}$T8I assembly EM density overlaid with the refined model. (b) B-factor distribution of the $Gag_{\Delta MA}$T8I assembly refined model. (c) Fourier Shell Correlation (FSC) of the CA-SP1 map from $Gag_{\Delta MA}$T8I assemblies (black), $Gag_{\Delta MA}$T8I assemblies with IP-6 (dark pink) and GagT8I VLPs (blue). (d) FSC between the CA-SP1 map and the refined model from $Gag_{\Delta MA}$T8I assemblies (black), $Gag_{\Delta MA}$T8I assemblies with IP-6 (dark pink) and GagT8I VLPs (blue).

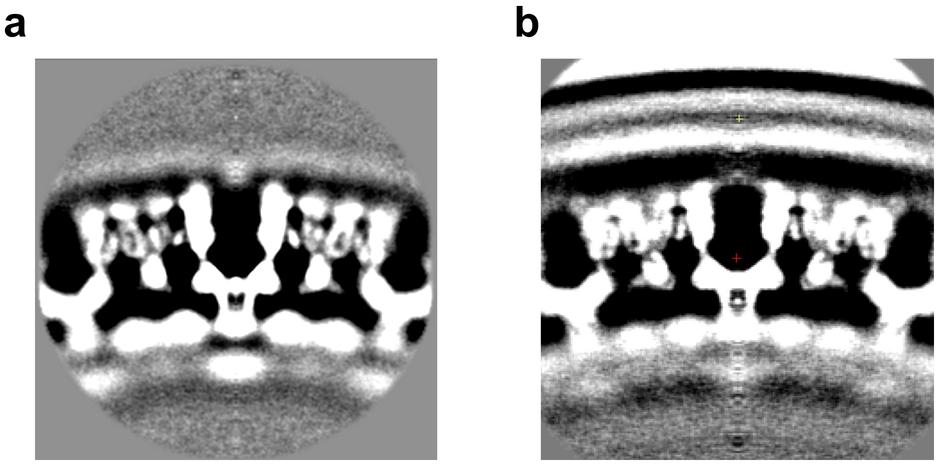

**Supplementary Figure 4 | GagT8I VLPs have diffuse density for MA.** (a) Gag$_{\Delta MA}$T8I unweighted and unmasked subtomogram average map. (b) GagT8I VLPs unweighted and unmasked subtomogram average map showing diffuse density for MA layer and lipid membrane.

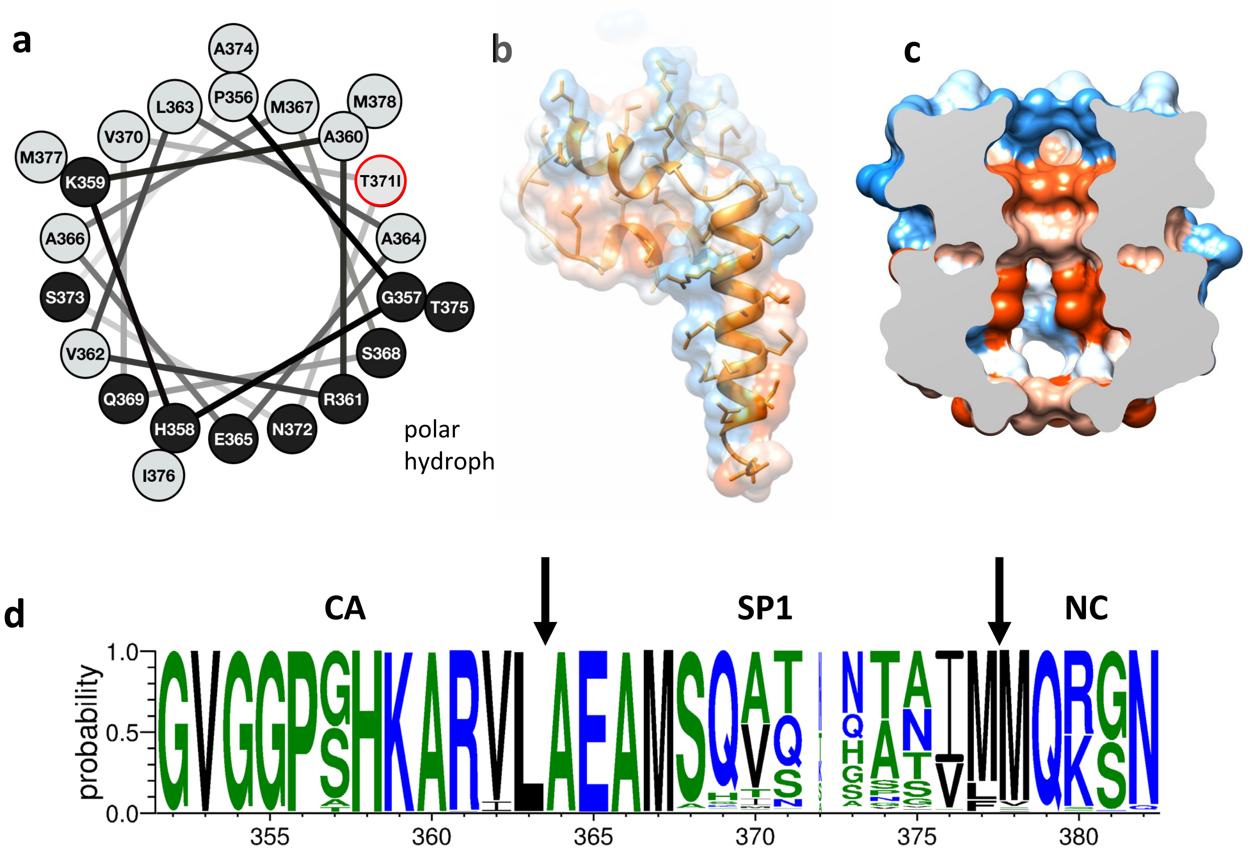


**Supplementary Figure 5 | The T8I mutation stabilizes the six-helix bundle by increasing its amphipathic nature.** (a) Helical wheel depiction of CA-SP1 residues, from P356 to M378. Polar residues are black, and nonpolar residues are gray. The T371I mutation is circled in red (modified from [12]). (b) Hydrophobicity surface of GagT8I from residue L290 to M377. Hydrophobic residues are colored in orange and polar residues in blue. (c) Hydrophobic residues are lining the inner surface of the six-helix bundle. (d) Sequence conservation of HIV Gag from G350 to N382 (produced with weblogo3 [13]).

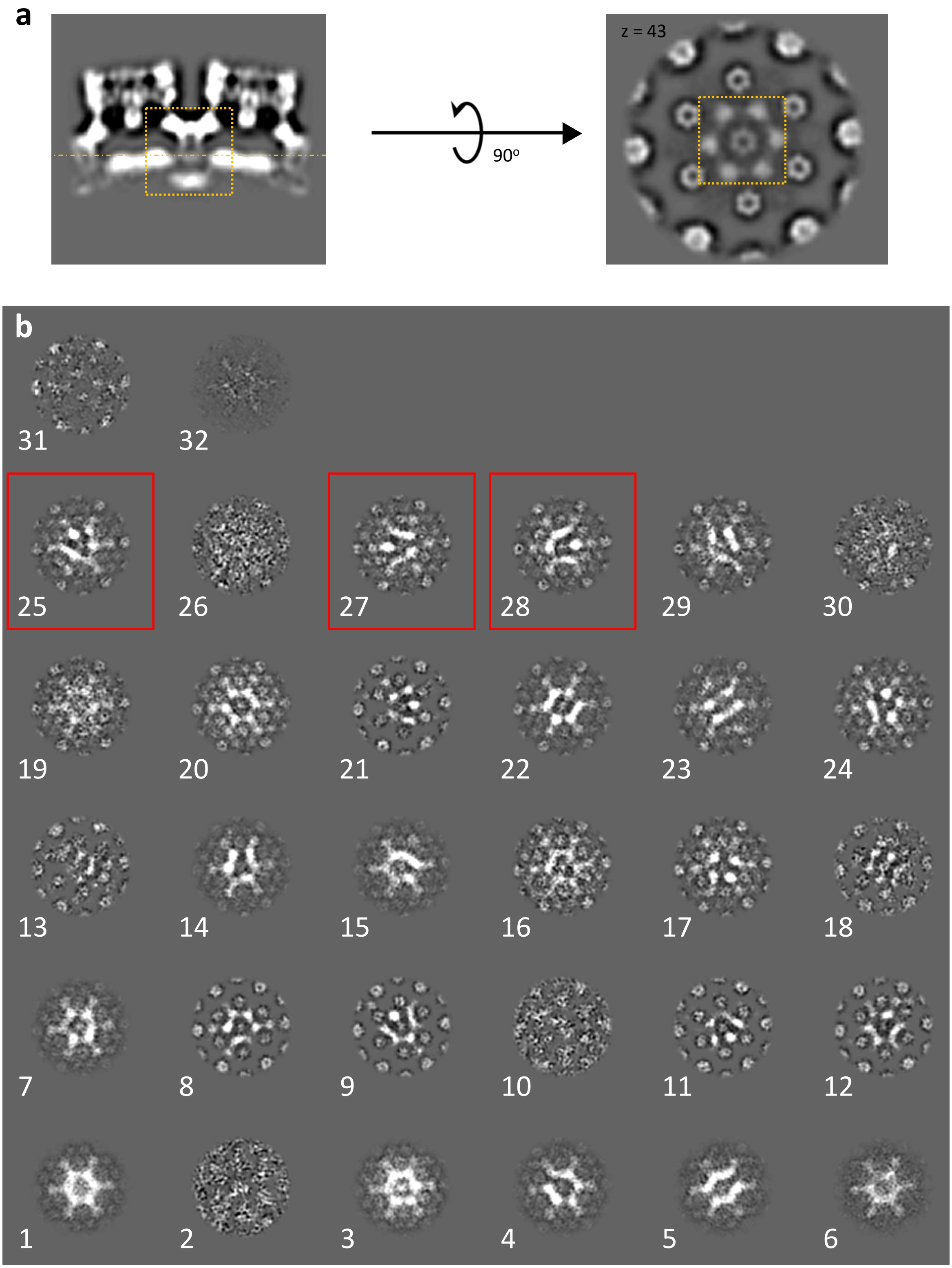


**Supplementary Figure 6 | Focused classification of NC-RNA density from Gag$_{\Delta MA}$T8I assemblies.** (a) Side (left) and top (right) views of CA-SP1 density map from Gag$_{\Delta MA}$T8I assemblies. The orange dashed box indicates the region used for classification. (c) 3D classes of NC-RNA layer. Classes in red squares, totalling 6747 subtomograms, were combined and averaged. No symmetry was imposed during classification and refinement.
